# Disruption by virtual reality of the cortical oscillations related to visuotactile integration during an embodiment process

**DOI:** 10.1101/2020.03.27.010942

**Authors:** Noriaki Kanayama, Masayuki Hara, Kenta Kimura

**Affiliations:** Human Informatics Research Institute, National Institute of Advanced Industrial Science and Technology (AIST), Tsukuba, Japan; Brain, Mind and KANSEI Sciences Research Center, Hiroshima University, Hiroshima, Japan; Graduate School of Science and Engineering, Saitama University, Saitama, Japan

**Author notes:** Corresponding Author Noriaki Kanayama, Human Informatics Research Institute National Institute of Advanced Industrial Science and Technology (AIST), Tsukuba Central 6, 1-1-1 Higashi, Tsukuba, Ibaraki 305-8566, Japan.

**Keywords:** virtual reality, rubber hand illusion, cortical oscillations, visuotactile integration, virtual reality noise, multisensory integration process

## Abstract

Virtual reality (VR) enables fast, free, and highly controllable experimental body image setting. Illusions pertaining to a body, like the rubber hand illusion (RHI), can be easily conducted in VR settings, and some phenomena, such as full-body illusions, are only realized in virtual environments. However, the multisensory integration process in VR is not yet fully understood, and we must clarify the limitations and whether specific phenomena can also occur in real life or only in VR settings. One useful investigative approach is measuring brain activities during a psychological experiment. Electroencephalography (EEG) oscillatory activities provide insight into the human multisensory integration process. Unfortunately, the data can be vulnerable to VR noise, which causes measurement and analytical difficulties for EEG data recorded in VR environments. Here, we took care to provide an experimental RHI setting using a head-mounted display, which provided a VR visual space and VR dummy hand along with EEG measurements. We compared EEG data taken in both real and VR environments and observed the gamma and theta band oscillatory activities. Ultimately, we saw statistically significant differences between congruent (RHI) and incongruent (not RHI) conditions in the real environment, which agrees with previous studies. No difference in the VR condition could be observed, suggesting that the VR setting itself altered the perceptual and sensory integration mechanisms. Thus, we must model this difference between real and VR settings whenever we use VR to investigate our bodily self-perception.

## Introduction

Many studies have demonstrated the successful embodiment of an external object that is not part of the real body since the rubber hand illusion (RHI) was first reported (Botvinick & Cohen, 1998). Generally, humans recognize external objects as their own body parts through a mechanism that connects real body parts and external objects in the form of dummy body parts based on multisensory (visuotactile) integration processing. A typical RHI is induced by synchronous, spatially congruent brush stroking stimuli on our occluded real hand, providing the tactile stroking information, and the visible dummy hand, providing visual stroking information. Theoretically, the multisensory integration of this visuotactile sensory information from the real and dummy hands enables the correspondence between the actual body and the external object. This results in the idea that “I felt as if the rubber hand was my hand,” termed RHI.

Interestingly, some prior art pieces have provided examples of RHI that could not be explained by this multisensory integration theory. For instance, asynchronous stimulation induced RHI in some participants who interpreted the this stimulation as synchronous (Costantini et al., 2016). Notably, this may be consistent with a common RHI research finding where some participants report asynchronous or incongruent sensations, but these results have not been formally reported because they are typically diminished by the averaging done across participants for published manuscripts. Visuotactile stimulation, with a slight delay on the visual stimulus (< 300 ms) has been shown to induce RHI (Shimada, Fukuda, & Hiraki, 2009) via multisensory integration. This suggests that asynchronous RHI can in fact be explained by the multisensory integration theory, as some participants with a broad multisensory integration window can effectively integrate asynchronous stimuli. Teramoto, Honda, Furuta, & Sekiyama (2017) demonstrated that aging can impact the visuotactile integration window for peri-personal space, indicating that multisensory integration mechanisms depend on the personal, physical, and/or psychological state of the body. Interestingly, Ferri et al. (2014) recently demonstrated that a tactile stimulus was not necessary for RHI, and that just an expectation of tactile stimuli can induce RHI. Thus, we must consider the possibility that the sensory integration itself is not needed for RHI, but rather just the prediction that visuotactile integration would occur.

The full-body illusion (FBI) is another example of the embodiment phenomenon. Although we generally consider this as a simple extension of RHI to the full body, it could pose a challenge to the multisensory integration theory for RHI. Specifically, FBI is a sense of “It felt as if the virtual body (or mannequin or object) in front of me was (part of) my body.” Participants must use a head-mounted display (HMD) and see the back image of themselves to induce this illusory experience, and synchronous brush stroking on the back or tapping on the collarbone can then induce the illusion (Ehrsson, Holmes, & Passingham, 2005; Lenggenhager, Tadi, Metzinger, & Blanke, 2007). For RHI, multisensory integration was restricted by the spatial congruence between real and dummy hands, within the extent of ecological validity. For example, a rubber hand that was rotated 180° could not induce RHI (Pavani, Spence, & Driver, 2000; Spence, Pavani, & Driver, 2004). Contrarily, FBI violates this spatial rule, because the participant’s body in the HMD is in front of the actual participant, which contradicts ecological validity. Thus, wearing an HMD to experience VR may change and expand the current multisensory integration theory.

Evans and Blanke (2013) demonstrated that illusions such as RHI could be achieved with a virtual hand while wearing an HMD, suggesting that RHI is not impacted by the environment, even when there is no direct comparison of actual reality and VR. In this case, they measured electroencephalography (EEG) activity during RHI induction and reported that the alpha/mu activation was differentiated by an ownership illusion, whether the visuotactile stimulations were synchronous or asynchronous. Notably, there was no significant effect on the gamma band activity, implying that gamma band oscillatory activity related to multisensory integration may be diminished in VR. All EEG activity in the study was obtained via power spectral density for a 2-second stimulation period. Interestingly, the gamma band oscillatory activity related to the visuotactile integration was very short-lived, lasting less than 100 ms, and its response could therefore be missed by the power spectral density for a 2-second stimulation. Therefore, to add clarity, we used two environments, real and VR, to induce RHI in a single participant and compared the resulting subjective ratings and EEG data.

Unfortunately, EEG studies in VR are prone to noise contamination. The VR environment offers experimental settings that allow the body representation to be manipulated quickly, freely, and controllably (Spanlang et al., 2014). However, when we also want to measure EEG, noise sources can harm the quality of the EEG data recorded. Contrarily, a recent Event-related potential (ERP) study did not report any significant noise contamination that could spoil the conditional differences between ERP measurements (Harjunen, Ahmed, Jacucci, Ravaja, & Spapé, 2017). Generally, oscillatory activities at high frequency band were more vulnerable to noise contamination than this ERP. Additionally, positive and negative deflections, at the exact opposite phases, were zero after averaging. The jittered strong deflection on the waveforms could be averaged out as average ERP, whereas the event-related spectrum powers (ERSPs) inherently included the power increase (Tallon-Baudry & Bertrand, 1999). One of the target components in this work is the gamma band oscillatory activity observed during visuotactile integration, which is both short-lived and vulnerable to noise contamination (Kanayama, Sato, & Ohira, 2007a). Therefore, we first tried to capture a less robust activity, using HMD, and clarify the difference in the visuotactile integration process between the real and VR environments. This could shed light on the variance of multisensory integration process depending on the circumstance. If we have any difference on EEG oscillatory activities related to multisensory integration process between real and VR, we could get any insights on the full body illusion seemingly violating the multisensory integration rule. If we have completely same properties between these environments, referring this fundamental knowledge, we can go on the study freely modifying our body in VR.

## Methods

### Participants

Thirty-two healthy individuals participated in this experiment. A participation fee was paid based on the National Institute of Advanced Industrial Science and Technology (AIST) guidelines. The average age was 23.38 ± 4.05, and the full age range extended from 20 to 40 years. Half of the participants were female, and half were male. All participants had normal or corrected-to-normal vision, and none reported neurological or psychiatric problems. The experimental procedures were approved by the ethical committee of AIST. All participants understood the details of the experiment prior to their participation, and written informed consent was obtained from each participant prior to the experiment.

### Materials and Apparatus

We modified the typical RHI experimental settings for our EEG measurement (Hiramoto et al., 2017; Kanayama et al., 2007a). A dummy rubber hand was made by filling a light-yellow kitchen glove with cotton. The natural posture of a left hand on a desk was mimicked using aluminum wire. Two white LEDs were attached to the tips of the index and ring fingers of the rubber hand. Participants wore an identical kitchen glove on their own left hand. Tiny cuts were made at the tips of the index and ring finger of the kitchen glove and bone-conducting earphones were inserted to make direct contact between the earphones, as a vibrator, and the finger skin surface. A 50 ms, 100 Hz sine wave was used to vibrate the bone-conducting earphones, and the intensity of this vibration was adjusted using a microphone pre-amplifier (QuadMic II, RME audio, Haimhausen, GERMANY). A cardboard box was used to occlude the participant’s real hand.

An HMD (HTC VIVE) was used for participants to see the VR environment. The environment of the experimental room, including LED-like spheres, was established in the software platform Unity 3D 4.5.4 platform (Unity Technologies, San Francisco, CA). The LED light in the VR environment was implemented in Unity using Emission parameters. The rigged pepper full hand, which is a hand model looks similar to the kitchen glove filled by cotton, in the assets of Leap Motion was used to represent the rubber hand in the VR environment. Stimulus timing was controlled by a custom C# algorithm. The concurrent presentation of the visual and auditory stimuli, as a stand-in for tactile stimulus, occurred in two successive lines. The program had an oscilloscope-confirmed 140 ms discrepancy (PicoScope 2204A, Pico Technology, Cambridgeshire, UK), which was kept in place to synchronize the presentation.

The Transistor-Transistor Logic (TTL) signal as a trigger was sent to the EEG amplifier simultaneously with the stimulus presentation through the parallel port. The HMD fixation belt was displaced to avoid any movement noise. Specifically, we hung the HMD on a hook from the ceiling and fixed it to each participant’s face with a Velcro belt at the neck, which allowed the movement to be noise-free during EEG measurements.

The power supply of the tracking camera was a mobile battery (iMuto M5). The DC power supply for the VIVE tracking camera was used without a ground in the shield room to help avoid noise from the commercial power supply. Participants wore an earphone to hear white noise (60 dB) to mask any auditory stimuli that occurred during the experiment. The response time was recorded with a custom recording device featuring a photosensor (BP-240-1001, Brain Products GmbH), which enabled us to precisely determine the onset of the finger movement. Figure 1 shows the experimental setup.

**Figure 1.**
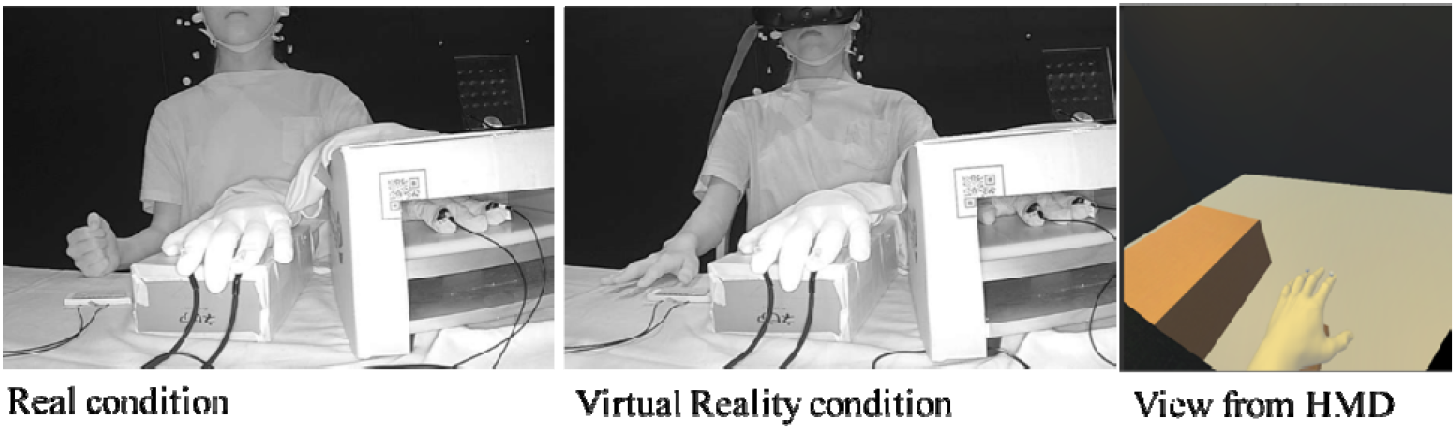
Experimental settings of RHI in both real and VR environments. In the left frame participants viewed the real visual space, in which the real kitchen glove was located in front of the participant while a cardboard box and black cloth occluded the participant’s left arm and hand. In the middle frame, the participant viewed the VR replication (shown in the right frame of the visual environment through the HMD.

### Experimental design and procedures

Our experiments focused on two within-participant factors: the real and VR environments, and either congruency or incongruency. The experiment was divided into two sessions for environmental modulation. Participants experienced visuotactile stimulation to from LEDs and vibrators to induce RHI in the real environment (Kanayama et al., 2007a; Kanayama, Sato, & Ohira, 2009), while they wore the HMD to experience the VR version of the experimental setup. Except for the HMD, all other settings were consistent between the real and VR environments. Further, the order in which the two environments were experienced was counterbalanced across all participants.

In each session, participants performed tasks for eight blocks, described in detail below. The first and last two blocks provided control conditions. In these, participants received only one sensory input, either LED light as a visual stimulus or vibration as a tactile stimulus, and were confirmed to not experience the RHI. The four blocks in the middle of the session featured either congruent or incongruent multisensory conditions, the order of which was counter-balanced. Participants received simultaneous visuotactile stimulation during each trial. In the congruent condition, the location of the visuotactile stimulation was consistent to induce visuotactile biding for RHI. For example, the LED emitted at the index finger of the rubber hand and the vibrator was felt at the index finger of the real hand. In the incongruent condition, the location of the visuotactile stimulation differed, to disturb visuotactile biding for RHI. For example, the LED emitted at the index finger of the rubber hand while the vibrator was felt at the ring finger of the real hand.

The experiments were broken into blocks, each of which consisted of 60 trials. The stimulated finger was pseudorandomly chosen. Each block began with a blank condition, defined as a 2 s period without stimulation. Participants then received 50 ms of visuotactile stimulation and were required to indicate which finger had been stimulated, as quickly as possible, by moving the finger on their right hand that corresponded to the stimulated finger on their left. Stimulus onset asynchrony (SOA) between the two trials was 1600, 1650, 1700, or 1750, and was pseudo-randomized for each trial.

After the 60 trials, participants were required to answer questionnaires on their subjective feeling of RHI, by stating a number from 0 to 100. The following items were included in the survey: (1) “I felt as if the rubber hand was my hand,” to illustrate the body ownership illusion intensity during each of the 60 stimulations, (2) “I felt as if the vibration was felt at the rubber hand,” to determine the touch referral illusion, (3) “I felt as if the LED emitted on my hand,” to note the spatial confusion of the visual event, (4) “I felt as if my hand was located where the rubber hand was,” to understand the subjective feeling of proprioceptive drift, (5) “I felt as if my hand was turning ‘‘rubbery.’’,” as a control question.

Immediately following these subjective reports, participants performed the proprioceptive drift task. They closed their eyes and pointed out the location of “the tip of the index finger of their own left hand” using the index finger of their right hand. At the beginning of this task, participants were asked to raise their arm to shoulder level avoiding touching the cardboard box which occluded their real hand during the task. Participants were then required to lower their right hand and move it horizontally to the left. When they found “the tip of the index finger of their own left hand,” participants were asked to say “here” and maintain that posture. At that time, the experimenter recorded the distance between the actual and indicated location of “the tip of the index finger of their own left hand.” Participants were instructed to ignore the vertical and depth directions during this task and focus only on the horizontal position. After the proprioceptive drift task and a short break, the next block started. The overall experimental flow is summarized in Figure 2.

**Figure 2.**
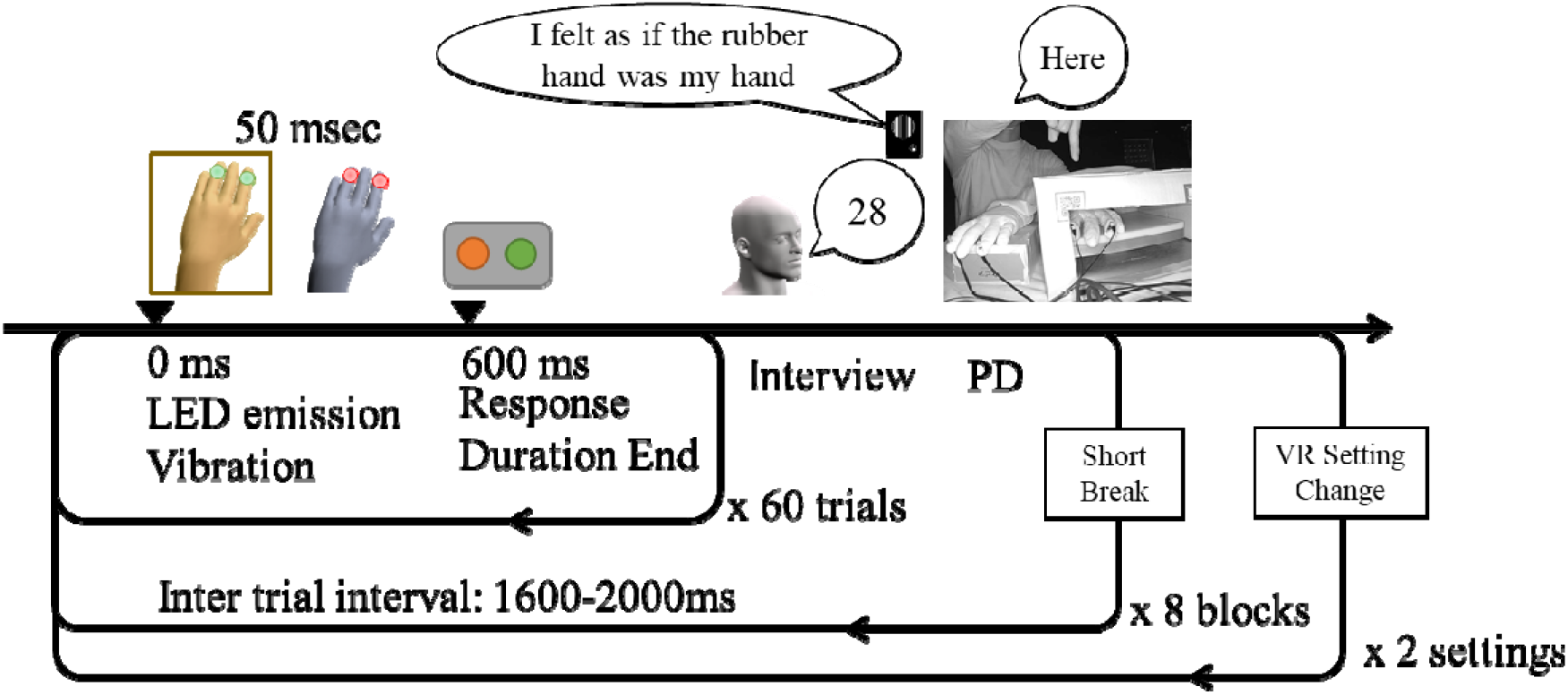
Experimental flowchart. Note that 0 ms indicates the stimulus onset of a single trial and the zero point of the time course for EEG segmentation. The abbreviation PD represent proprioceptive drift, wherein participants pointed the tip of their left index finger withou seeing it.

### Behavioral data and analyses

We obtained the reaction times and the rate of correct responses for stimulated finger detection in each trial. These data were averaged across trials, using one value for each condition and each participant. Subjective ratings for the five questionnaire items and a single proprioceptive drift value were also obtained for each condition and each participant. Paired *t*-tests were conducted to determine the difference between the congruent and incongruent conditions in the same environment, and the difference between environments for the same stimulation condition. The *p*-value was corrected using Bonferroni methods.

### EEG measurement and analyses

Continuous EEG waveforms were obtained for each condition and each participant, using an actiCHamp amplifier (Brainproducts GmbH, Munich, Germany), from 63 electrodes distributed over the scalp. Electrode locations were selected based on the 10-10 international standard EEG electrode placement, using an EasyCap (EasyCap, GmbH, Herrsching, Germany). A reference electrode was located at the Cz, and the grand electrode was located at the Fpz. The sampling rate was 1000 Hz. The recorded waveforms were filtered with the hardware filter, and the low-frequency cutoff was 0.016 Hz with a time constant of 10 s. The impedance at each electrode was maintained at least below 30 kΩ, and typically below 10 kΩ.

Subsequent data analyses were conducted using EEGLAB, version 14_1_1b (Delorme & Makeig, 2004; Delorme et al., 2011) under Matlab R2018a (MathWorks, Natick, MA, USA). A digital high pass filter, at 1 Hz, was first applied to the recorded waveforms, and the Clean-Line plug-in (Mitra & Bokil, 2008) was conducted at 50 Hz and 100 Hz to reduce noise from the commercial power supply. The cleaned waveforms were then segmented into the corresponding trials. The zero-time point of each segment was defined as the visuotactile stimulation onset (0 ms), and segments ranged from −600 to 1200 ms, relative to the stimulus onset. The first independent component analysis (ICA) was conducted on segmented data and captured 63 independent components (ICs) for each participant. Artifact-contaminated trials were detected using certain indices calculated from the IC waveforms: maximum and minimum amplitude, mean trial probability, kurtosis value, and spectrum power. The artifact criteria for individuals was adjusted to keep the total discarded trials below 10%. Using the datasets after the artifact trial rejection, the second ICA was conducted to obtain 63 new ICs for each participant. The dipole estimation was conducted on these ICs using dipfit2 (EEGLAB plug-in using FieldTrip toolbox functions; Oostenveld et al., 2011). Thus, the total number of ICs was 2016 (32 participants x 63 ICs).

Among all the obtained ICs, cortical activation-related ICs were selected based on the residual variance of the dipole estimation (< 15%). The remaining 686 ICs were clustered using the k-means method. We used the scalp topography and dipole location for this clustering. The dimension of the scalp topography data was reduced to ten by principal component analysis. The clustering number of 13 was determined based on the Davies Bouldin criteria.

The ERSPs, during 0-800 ms post-stimulus and 3-90 Hz, were calculated using a Morlet wavelet (cycles at 3 and 90 Hz are 2 and 12, respectively). Additionally, ERSP during 300-50 ms pre-stimulus was calculated as a baseline, which was used to convert the post-stimulus data to dB. The statistical difference in the ERSPs between conditions was tested using Monte Carlo permutation statistics, with a cluster correction. The triangulation method was used to make the channel neighbor, and the MaxSum method was used for clustering (Cohen, 2014) and implemented in FieldTrip (corrected p < 0.05).

## Results

### Subjective Feeling of RHI

To start, we compared the subjective ratings data for each condition (Figure 3) and tested the statistical difference of the subjective RHI experience using 2 factor ANOVA to confirm that RHI was successfully induced. We found a significant environmental effect, where real was less than VR, for Q1 (F (1,31) = 23.51, p < 0.01, eta = 0.17), Q2 (F (1,31) = 11.93, p < 0.01, eta = 0.13), Q3 (F (1,31) = 12.23, p < 0.01, eta = 0.12), Q4 (F (1,31) = 27.17, p < 0.01, eta < 0.23), and Q5 (F (1,31) = 9.30, p < 0.01, eta < 0.05).

**Figure 3.**
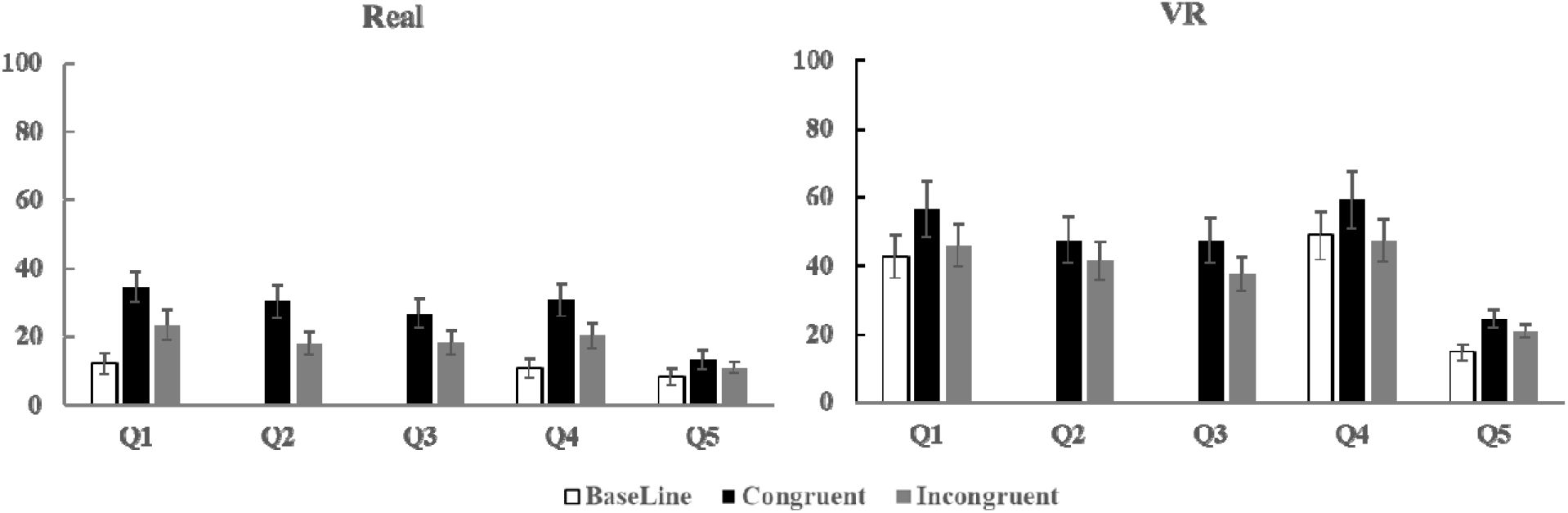
Averaged RHI subjective ratings and standard-errors, where participants wer required to state a number between 0-100, with 0 and 100 indicating “Did not feel at all” an “Felt very strongly,” respectively, to describe the extent to which they agreed with each state feeling. Q1 was “I felt as if the rubber hand was my hand”, Q2 was “I felt as if the vibration wa felt at the rubber hand”, Q3 was “I felt as if the LED emitted at my hand”, Q4 was “I felt as i my hand was located where the rubber hand was”, and Q5 was “I felt as if my hand was clad I rubber”. In the legend, “Real” and “VR” refer to the real and VR environment sessions respectively. The “Baseline” represents ratings given before the experimental session began. “Congruent” and “Incongruent” represent ratings obtained after stimulations in the congruen and incongruent conditions, respectively.

We also found a significant differences between baseline, congruent, and incongruent conditions for Q1 (F (2,62) = 21.60, p < 0.01, eta = 0.06), Q4 (F (2,62) = 18.61, p < 0.01, eta < 0.04), and Q5 (F (1,31) = 4.72, p < 0.05, eta < 0.02). Furthermore, we found significant differences between congruent and incongruent condition items for Q2 (F (1,31) = 10.93, p < 0.01, eta = 0.03) and Q3 (F (1,31) = 18.61, p < 0.01, eta = 0.03).

Significant effects for the proprioceptive drift were similar (environment, F (1,31) = 20.30, p < 0.01, eta = 0.08; condition, (F (2,62) = 14.55, p < 0.01, eta = 0.06); no interaction).

### Behavior results for tactile detection task

Next, we calculated the inverse efficiency (Figure 4) as the reaction time (ms) divided by the correct response rate (%) to quantitatively evaluate the tactile detection task performance. Specifically, a higher inverse efficiency corresponds to a worse performance. Here, we used paired t-tests to find any significant differences between the tested conditions and environments. In both real and VR environments, the differences between congruent and incongruent conditions were significant (real: t (31) = −3.12, p < 0.05 [0.003821], VR: t (31) = −4.61, p < 0.01 [0.0006584]). However, a significant difference between real and VR environments was only observed under the incongruent condition (t (31) = −2.89, p < 0.05 [0.007004]). The congruency effect, calculated by subtracting the inverse efficiency of congruent and incongruent conditions, is used as an index of multisensory binding. In this case, we did not observe a significant difference between the real and VR environments (107.40 at Real vs. 149.04 at VR, t (31) = 0.94, n.s.).

**Figure 4.**
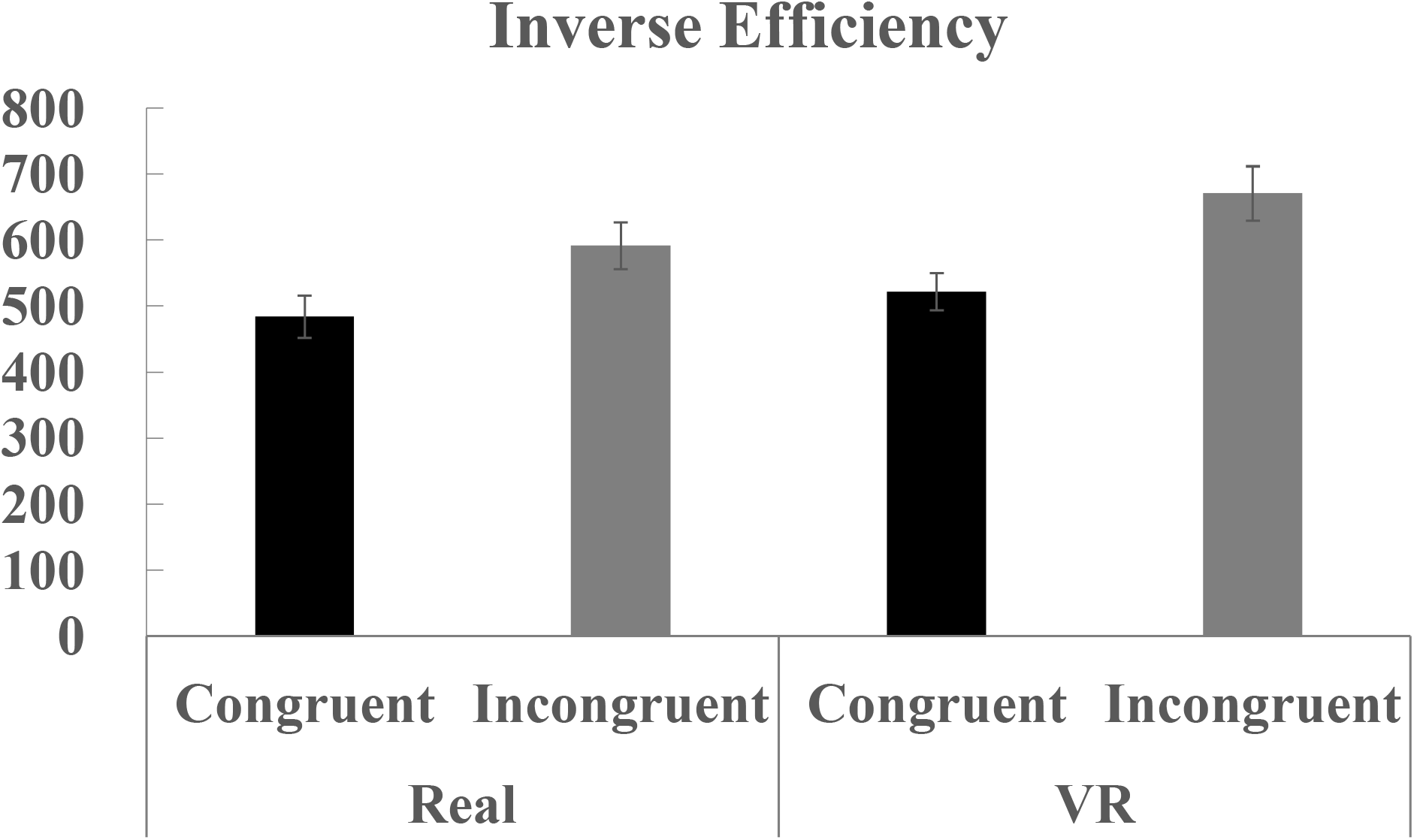
Average inverse efficiency for the tactile detection task on either the index or ring finger. Error bars indicates standard errors. Participants responded with which finger was stimulated. “Real” and “VR” indicate results for real and VR environment sessions, respectively. “Congruent” and “Incongruent” represent scores rated after stimulations done under congruent and incongruent conditions, respectively.

### EEG results

When evaluating the EEG data, ICA clustering analysis revealed 13 clusters of cortical activity, summarized in Table 1, that are related to the visuotactile integration process. In this work, we focused on the parietal cluster and the precentral (premotor) cluster, based on previous reports (Kanayama et al., 2007; 2016), and ultimately targeted clusters 6 and 10 as parietal and precentral clusters, respectively. Both of these showed low residual variances when averaged across all components involved in the cluster (Cls6: 3.95 %, Cls10: 4.66 %).

**Table 1.**
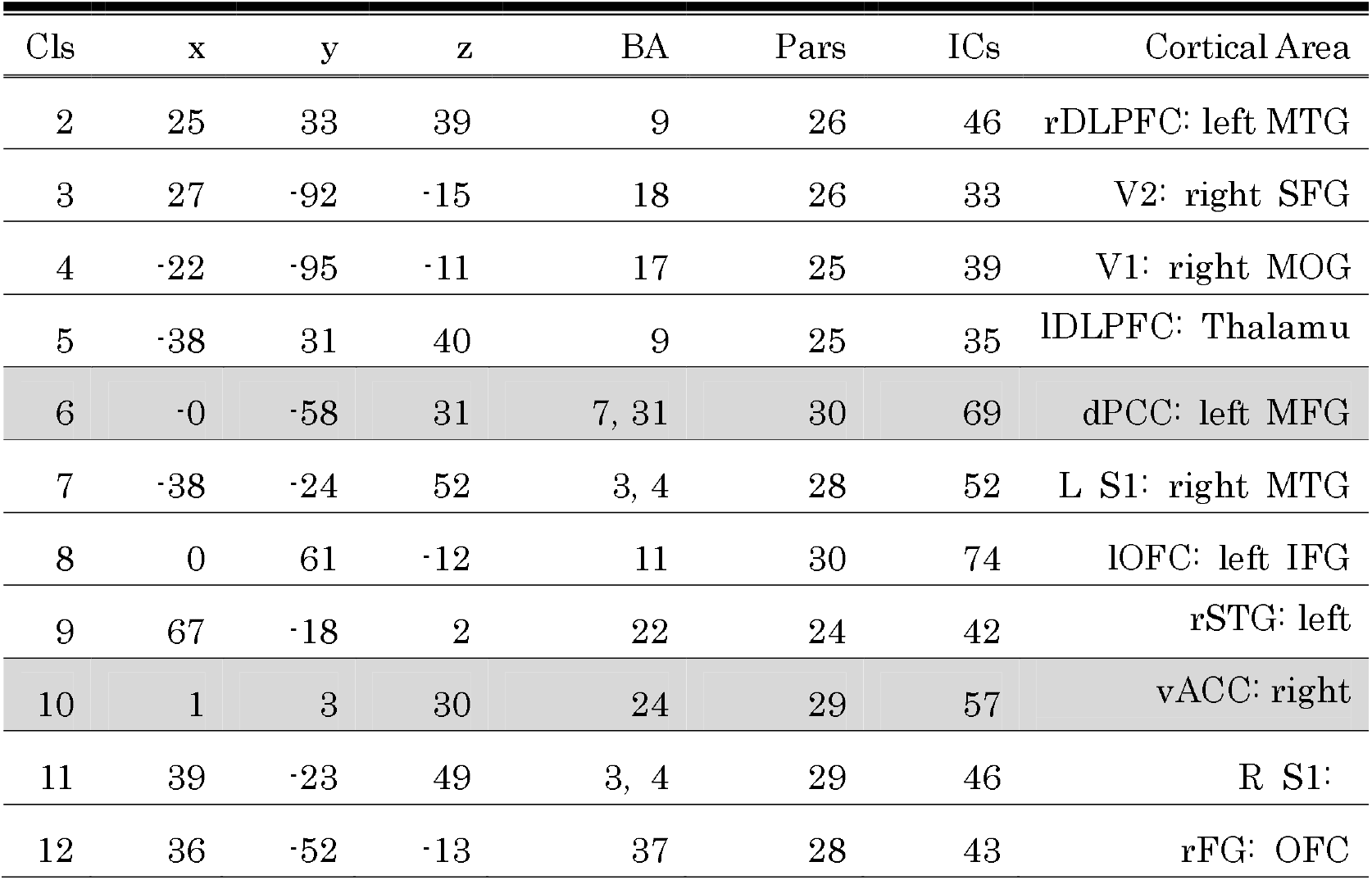
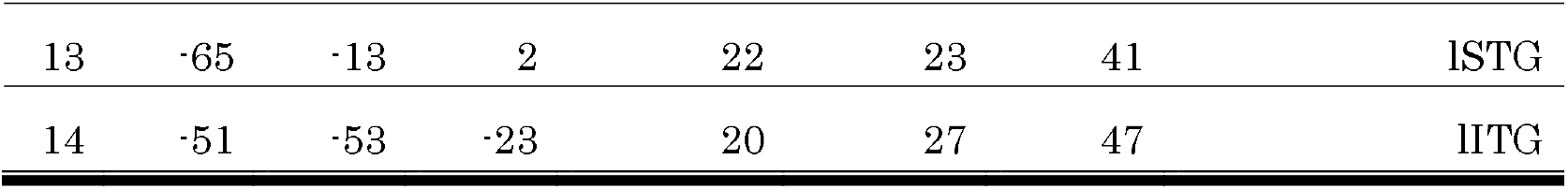
MNI coordinates of the averaged dipole location for all clusters. Cluster number 1, the parent clusterwas excluded. The shaded rows correspond to the clusters selected for future analyses. The Brodmann Area (BA) was detected using the Talairach coordinates, which were converted from the MNI coordinates. Cls: cluster, Pars: number of participants, ICs: the number of Independent Component included in the cluster. MTG: Middle Temporal Gyrus, SFG: Superior Frontal Gyrus, MOG: Middle Occipital Gyrus, MFG: Middle Frontal Gyrus, MTG: Middle Temporal Gyrus, IFG: Inferior Frontal Gyrus, OFC: Orbitofrontal cortex, SOM: Somatosensory Area, STG: Superior Temporal Gyrus.

To statistically compare the ERSPs between congruent and incongruent conditions, we separated the data into two frequency bands at high (30-90 Hz) and low (3-12 Hz) frequencies, denoted as the gamma and the low frequency (including theta) bands, respectively. We then focused on the target time window, 200-600 ms, for the high-frequency data, while 0-600 ms for the theta band data. Both the gamma band activity (Figure 5) at the parietal cluster and the theta band activity (Figure 6) at the precentral cluster were differentiated by congruent visuotactile stimulus in the real environment, while there was nothing in the VR environment. Two theta components separated in time, with the same statistical results, were found at about 50-300 ms and 400-800 ms.

**Figure 5.**
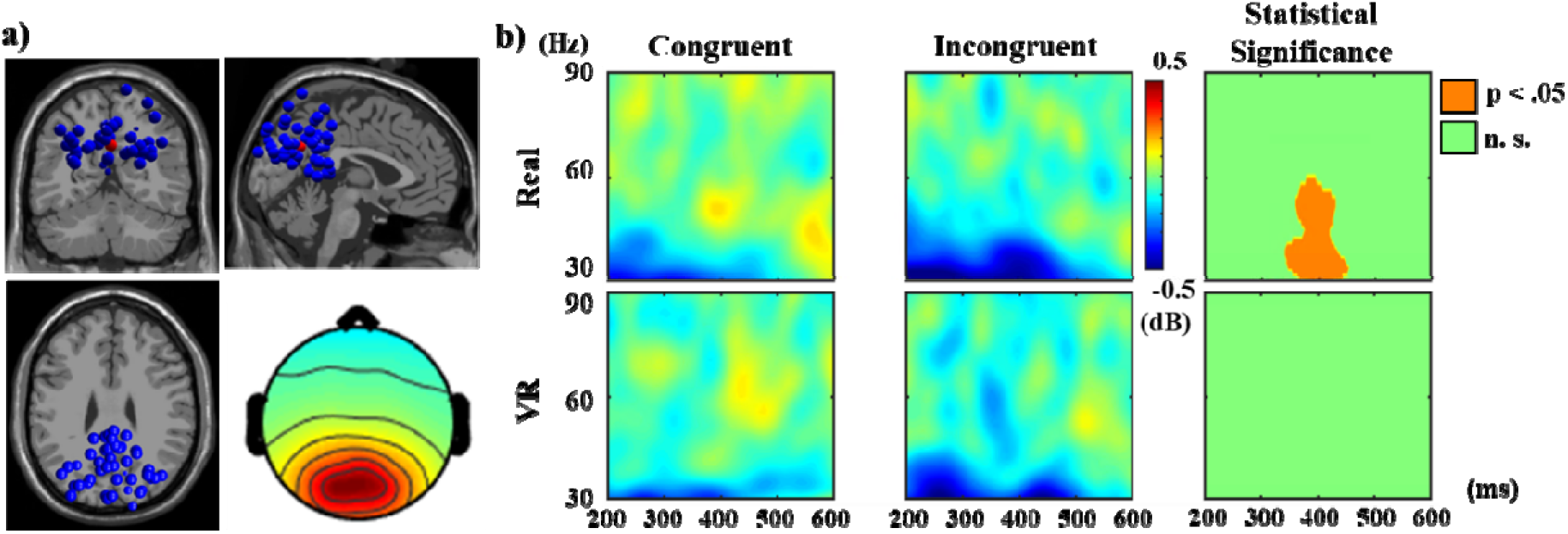
a) Dipole locations of all components (blue) involved in the parietal cluster and the centroid of all dipole positions (red) and the topographical map on the scalp averaged across all components. b) ERSPs at the gamma band (30-90 Hz) for each condition, averaged across all IC obtained from all participants involved in the parietal cluster (left and middle columns) and the statistical tests for the congruent condition (right column). The upper and lower rows provide the ERSPs obtained in the real and VR environment sessions, respectively.

**Figure 6.**
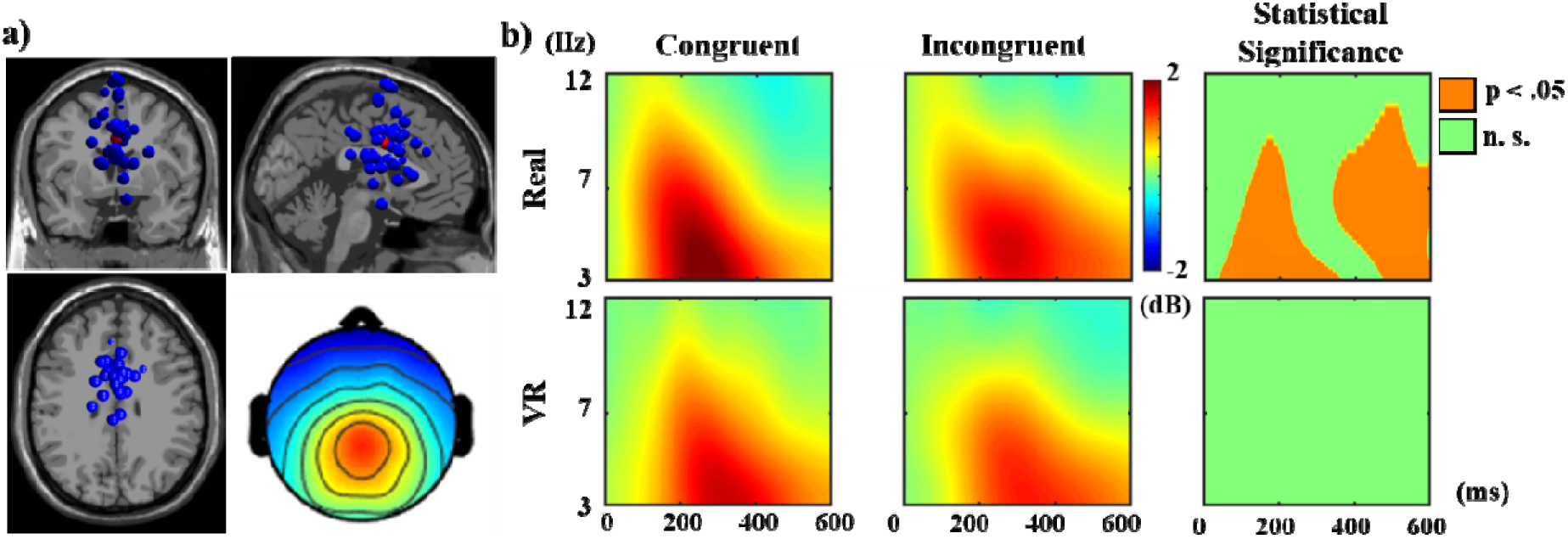
a) Dipole locations of all components (blue) involved in the precentral cluster and th centroid of all dipole positions (red) and the topographical map on the scalp averaged across al components. b) ERSPs at the theta and alpha bands (3-12 Hz) for each condition, average across all ICs obtained from all participants involved in the parietal cluster (left and middle columns) and the statistical tests for both congruent and incongruent conditions (right column). The upper and lower rows provide the ERSPs obtained in the real and VR environment session, respectively.

## Discussion

In this work, we investigated the multisensory integration process during RHI induction by evaluating EEG oscillatory activities. The specific Importantly, the EEG oscillations that relate to this process are different in a VR environment (Kanayama et al., 2007; 2008; 2016). Specifically, gamma and theta band oscillation related to sensory binding is observed during RHI in a real environment, but were diminished in VR. Clearly, our visuotactile sensory integration process is altered when we an HMD is worn to see the VR environment, and the RHI can in fact be induced by mechanisms other than multisensory binding. Thus, this work provides the first report indicating that a VR environment can disrupt the typical sensory integration process.

The subjective evaluation of RHI intensity revealed some similar and different points between real and VR environments. In the real environment, our experimental settings are similar to previous RHI experiments (Botvinick & Cohen, 1998). In the original RHI work and subsequent studies that replicated the illusory experience induced by synchronous visuotactile stimulation, a significant difference of subjective RHI intensity is expected between synchronous and asynchronous stroking. In our experimental paradigm, the synchronous and asynchronous comparison corresponded to the spatially congruent or incongruent, respectively, visuotactile stimulation using an LED and vibrator. This type of RHI induction has previously successfully shown a significant subjective RHI intensity difference between congruent and incongruent conditions (Pavani et al., 2000). Thus, our demonstration of the statistically significant condition dependence of the subjective RHI feeling, in a real environment, is consistent with previous works. Further, this condition dependence was replicated in the VR environment, suggesting that VR can induce similar subjective feelings of RHI that depend on whether the visuotactile stimulation is congruent or incongruent=. Similarly, illusory feelings resulting from RHI induction have previously been replicated in VR (Evans & Blanke, 2013).

Notably, there is a significant difference in the subjective RHI feeling between real and VR environments, which demonstrate that the RHI intensity was enhanced by VR environment. Interestingly, this difference was significant for the congruent stimulation condition, the incongruent stimulation condition, and even for the baseline condition. During the baseline period, participants had no experience with any stimulation of either visual and tactile sensory modality, but merely saw the VR environment mimicking reality. Specifically, the averaged subjective RHI values of Items 1 and 4 were slightly higher at the baseline period of VR sessions compared to the values obtained after congruent visuotactile stimulation in real sessions. Simply seeing the rubber hand in VR space can induce a stronger RHI experience, compared to the classical RHI, which cannot be attributed to visuotactile multisensory integration.

Contrary to the subjective rating data, the behavior index data obtained in the tactile detection task suggests that the results do not differ widely between the evaluated environments. Importantly, the congruency effect, which provides as a possible index of RHI (Hara et al., 2016; Pavani et al., 2000), was not significantly affected by the environment. Further, the behavioral response as a result of the multisensory integration process was not altered by VR, but we cannot confirm whether various internal processes, which all have the same observable outputs, were necessarily the same using these behavioral results.

Considering the EEG components, we can draw several conclusions about the altered sensory properties in VR. First, the increased gamma band oscillation under congruent conditions, compared to incongruent, was observed in the real environment but not in the VR environment. Generally, the gamma band oscillation was considered to have a role of information binding (Singer, 1999; Singer & Gray, 1995; Singer, 2001), and has been shown in the RHI paradigm to induce multisensory integration (Kanayama et al., 2007a). A mouse model using intracranial EEG revealed that gamma band oscillation could be detected during the sensory binding or visuotactile integration process (Quinn et al., 2014). Notably, we could not bind the tactile stimulation on our real hand and the visual inputs on the dummy hand. Second, theta band oscillatory responses increased significantly in the incongruent condition compared to congruent, which is consistent with a previous study investigating EEG oscillatory activities during visuotactile interactions (Kanayama & Ohira, 2009).

Additionally, the gamma band oscillation phase might become coupled to the theta band oscillations (Canolty et al., 2006). Bimodal peaks of theta activities around 200 ms and 400 ms, observed both before and after the gamma band oscillatory activity under congruent conditions, could be closely related to the multisensory integration process (Kanayama & Ohira, 2009). Contrarily, the theta band oscillations at the frontal area corresponds to the error-related response (Keil, Weisz, Paul-Jordanov, & Wienbruch, 2010; Luu, Tucker, & Makeig, 2004; Roger, Bénar, Vidal, Hasbroucq, & Burle, 2010; Trujillo & Allen, 2007; Yeung, Bogacz, Holroyd, Nieuwenhuis, & Cohen, 2007) and cognitive load and control (Cavanagh, Zambrano-Vazquez, & Allen, 2012; Cummins, Broughton, & Finnigan, 2008; Nigbur, Ivanova, & Stürmer, 2011). Thus, the increase in theta power observed in this study was attributed to the cognitive load required to process the incongruent visuotactile stimuli. However, this increased activity at the theta band, in the incongruent condition, was diminished in the VR environment, suggesting that VR enables us to uncouple from visuotactile competition on our body.

In our experiment, we never showed the participants’ left hand in VR setting. Participants could not see their own real left hand in the HMD display, and there was no corresponding VR hand. It would have been straightforward to use any realistic human hand model and taken a training session to familiarize subjects with the contingent moving of the realistic VR hand using a motion capture system, such as leap motion. However, we did not pursue this, as we hypothesized that an experience without one’s own left hand could potentially diminish the induced visuotactile integration processing. Therefore, further experiments are needed to compare the EEG oscillatory activities with or without this adjusted experience with the left hand in the VR environment. If the EEG oscillation is the same with and without an adjusting phase on the VR hand, the VR environment itself will be shown to have a strong impact on the EEG components related to the body ownership illusion and visuotactile integration, regardless of the experience of seeing and moving one’s hand in VR.

Another explanation for the observed EEG difference between real and VR environments is the impact of wearing the HMD. Additional investigations may illuminate any differences between reality and video of real space through an HMD. To investigate this, cameras attached at the front of an HMD could capture video to see the real environment, like our eyes would, rather than VR, and can deliver almost identical visual inputs to participants with the only difference being whether an HMD was used. Some research on infants suggests that video learning is less effective for developing brains, because it is not reality, and the brain responses to performing actions were significantly different from responses to video of the same content (Shimada & Hiraki, 2006). According to these findings, there should be a noticeable difference between the video and the reality included in our results.

The general lack of condition dependence in our results could be attributed to noise contamination. Especially at high-frequency ranges, EEG oscillatory activity can be quite vulnerable to noise. Furthermore, there is evidence of possible contaminations to oscillatory activities in EEG recorded on the scalp. The first of these is movement noise. Especially, in the gamma band, noise can be induced by electrode movement; hence, we took intentional steps to eliminate this movement by removing the fixation belt of the HMD onto the scalp. While participants with small faces experienced contact between the HMD face cushion and the Fp line electrodes, these electrodes have only a negligible impact on the target components, as a result of ICA and clustering; hence, this did not impact our results.

## Limitations of the research

While we did not control for past experiences with VR, fortunately almost half of the participants had some VR experience, while the others had none, and the group was confirmed ex-post facto. For future research, more thorough and intentional randomization would require a larger sample population.

There may have been contamination of refresh rate noise from the HMD, which was attached to the face near the frontal site of the scalp. Although a cushion sponge was inserted between the face and the HMD, thereby possibly insulating the HMD-related noise from the EEG recording, refresh noise could still contaminate the EEG waveforms through the spatial conduction of electromagnetic noise. We focused on frequencies below 90 Hz, which is the same frequency of the monitor refresh, for high-frequency gamma band oscillatory activities. Despite this vulnerability, we can still assume that the results for frequency bands 30-60 Hz were not at all affected by the HMD refresh noise.

Although the visual appearance of the experimental setup, including the rubber hand, cardboard box, and black cloth in VR observed through the HMD, closely resembled the real one, the artificial setup could still be easily distinguished. It is, therefore, likely that those visual stimuli in VR were completely different from those in the real environment. Additionally, the view angle in VR was limited to the HMD specifications, and continuously included the blacked-out space in the peripheral visual field. We can therefore assume that this difference in visual perception caused some of the difference in the EEG activity between real and VR environments. Specifically, gamma band oscillation was not observed in the unimodal, visual-only condition in reality (Kanayama et al., 2007); hence, we can conclude that the target components in this study were not merely modulated by unimodal stimulation.

## Conclusion

In this work, we measured the EEG oscillatory activities during RHI induction in real and VR settings and investigated the difference of the visuotactile integration processes between these environments. We observed generally stronger RHI experiences in VR, but a significant increase for the congruent condition over both the baseline and incongruent conditions remained subjectively the same in both real and VR environments. The EEG components related to the visuotactile integration process were diminished in the VR environment, despite the subjective similarity of the illusion. Although we have to make any further investigation controlling the VR experience or any other possible EEG noise contamination, this report, at least, suggests that the perception and integration of visuotactile input is drastically different in real and VR environments. When we investigate the cortical substrate of body representation in VR, we should have any attention on that multisensory integration rule, which affects body related recognition, can be modified by environment including HMD, augmented reality or any other virtual environment.

## Acknowledgments

The authors would like to thank Moena Okibuchi, Tomoko Otomo, Chiaki Hisamatsu, Saori Matsumoto (AIST, Japan) for their assistance with conducting the experiments. We would also like to thank Professor Hiroaki Shigemasu (Kochi Technology University, Japan) for his advice regarding VR experiments. This work was supported by the Japan Society for the Promotion of Science KAKENHI, Research Fellowships for Young Scientists, grant number 16H05958 and by Grant-in-Aid for Scientific Research (A), grant number 19H00631. In addition, the Japan Science and Technology Agency’s Center of Innovation Program, grant number JPMJCE1311, partially supported this research.

## Authors’ contribution

All authors have read, discussed, and approved the manuscript for submission. N.K. designed the behavioral paradigms. M.H. designed and developed the VR settings. N.K. collected the data. N.K. and K.K. discussed the construction of the paper. N.K. wrote the paper.

